# LRET-derived HADDOCK structural models describe the conformational heterogeneity required for processivity of the Mre11-Rad50 DNA damage repair complex

**DOI:** 10.1101/2021.08.04.455035

**Authors:** Marella D. Canny, Michael P. Latham

## Abstract

The Mre11-Rad50-Nbs1 protein complex is one of the first responders to DNA double strand breaks. Studies have shown that the catalytic activities of the evolutionarily conserved Mre11-Rad50 (MR) core complex depend on an ATP-dependent global conformational change that takes the macromolecule from an open, extended structure in the absence of ATP to a closed, globular structure when ATP is bound. We have previously identified an additional ‘partially open’ conformation using Luminescence Resonance Energy Transfer (LRET) experiments. Here, a combination of LRET and the molecular docking program HADDOCK was used to further investigate this partially open state and identify three conformations of ATP-bound MR in solution: closed, partially open, and open, which are in addition to the extended, apo conformation. These models are supported with mutagenesis and SAXS data that corroborate the presence of these three states and suggest a mechanism for the processivity of the MR complex along the DNA.

## Introduction

Mre11-Rad50-Nbs1 (MRN) is an essential protein complex required for the repair of DNA double strand breaks (DSBs) (Paull, 2018; Syed and Tainer, 2018). This complex recognizes the broken DNA and begins processing the break via Mre11 exo- and endonuclease activities and Rad50 ATP binding and hydrolysis (Paull, 2018). Nbs1, found only in eukaryotes, further modulates MR activity and signals downstream repair effectors to the site of the break (Deshpande et al., 2016; Oh et al., 2016). If DNA DSBs are not repaired, the cell may undergo cell death via apoptosis, or, if the break is not repaired correctly, a loss of genetic information or gross chromosomal rearrangements can occur potentially resulting in immunodeficiencies and cancer (Ciccia and Elledge, 2010; Oh and Symington, 2018). Many mechanistic and structural studies have been performed on the evolutionarily conserved Mre11_2_-Rad50_2_ (MR) core complex from bacteria, archaebacteria, and eukaryotes and have shown that the complex undergoes a dramatic ATP-induced global conformational change that is required for its various functions. This two-state model, which originated from X-ray crystallographic studies (Lafrance-Vanasse et al., 2015; Lammens et al., 2011; Lim et al., 2011; Möckel et al., 2012), has MR transforming from an extended arms-open-wide conformation, where the two Rad50 nucleotide binding domains (NBDs) are far apart in space, to a ‘closed’ conformation that sandwiches two ATPs between the NBDs in a more compact, globular structure (Fig. 1A). In the closed conformation, the Mre11 nuclease active sites are occluded and Rad50 can bind DNA (Liu et al., 2016; Möckel et al., 2012; Rojowska et al., 2014; Schiller et al., 2012; Seifert et al., 2016). Once Rad50 hydrolyzes the bound ATPs, the complex returns to the extended conformation where DNA substrates can again access the Mre11 active sites. Critically, ATP binding and hydrolysis, and therefore the cycling between states, appears to be required for DNA unwinding(Cannon et al., 2013), processive Mre11 nuclease activity (Herdendorf et al., 2011; Paull and Gellert, 1998), and downstream signaling events through ATM kinase (Cassani et al., 2019; Lee et al., 2013).

**Fig. 1.**
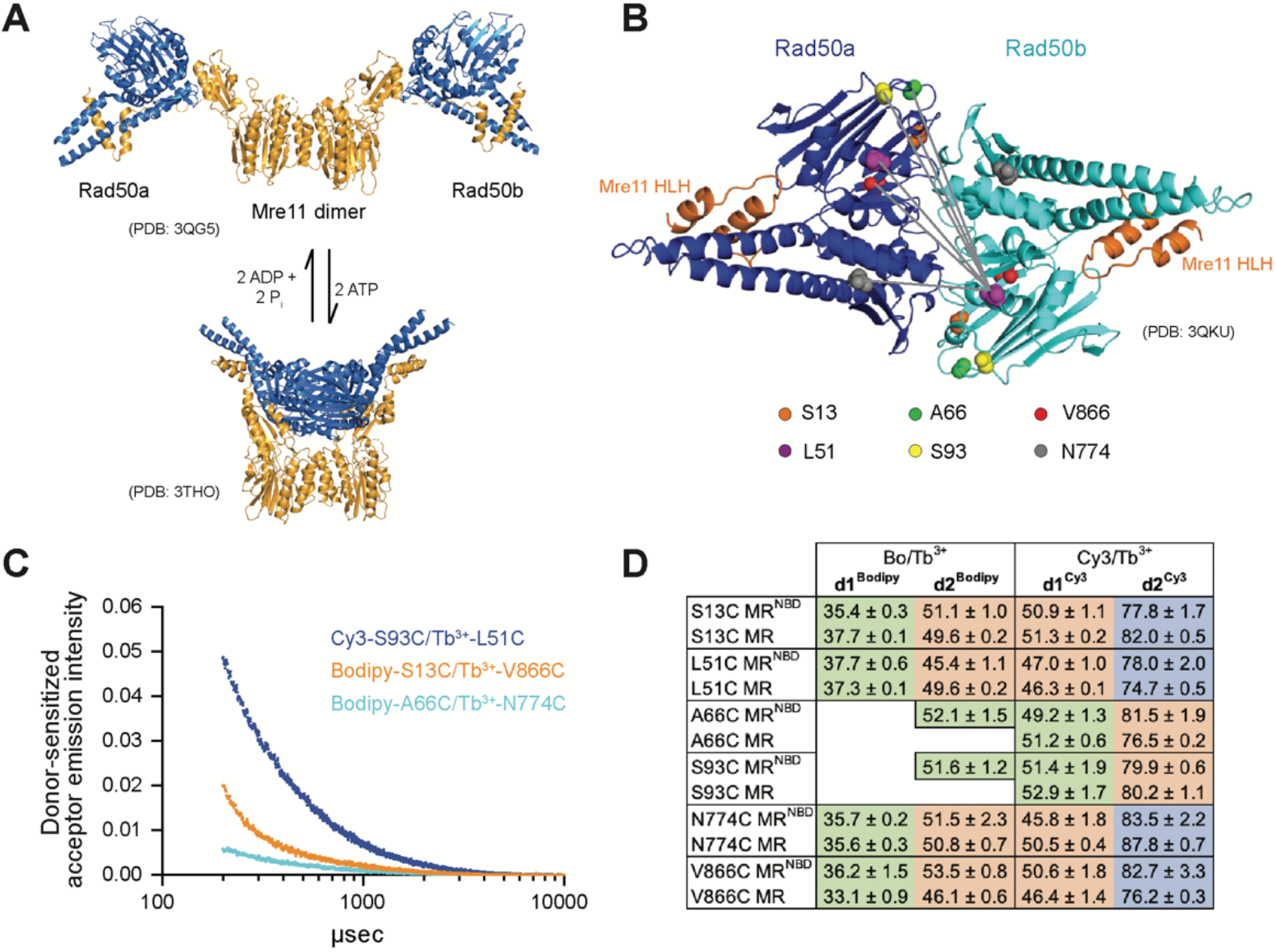
LRET measures distances between Rad50 residues in Pf MR^NBD^. (A) X-ray crystal structures of *T. maritima* MR^NBD^ showing the ATP-dependent transition between extended and closed conformations(Lammens et al., 2011; Möckel et al., 2012). (B) Positions of LRET probes highlighted on the *P. furiosus* Rad50 AMPPNP-bound dimer(Williams et al., 2011). Grey lines show L51 of Rad50b interacting with each of the probe residues of Rad50a. (C) Plot of representative LRET emission decays versus time after Tb^3+^-chelate donor excitation. (D) Table of the LRET-determined distances (in Å) for the identity pairs, where the same residue is labeled in each protomer, in MR^NBD^ and full-length MR complexes. One Rad50 was labeled with either with Bodipy FL or Cy3 acceptor and the other with Tb^3+^- chelate donor. Green, orange, and purple shaded cells indicate distances in the ‘closed,’ ‘partially open,’ and ‘open’ conformations, respectively. Values are the mean and standard deviation of at least three replicates.

Because the Mre11 active site is occluded in the closed conformation, it was hypothesized that MR nuclease activity originates from either the extended or an otherwise unknown intermediate structure (Deshpande et al., 2014; Lafrance-Vanasse et al., 2015; Lammens et al., 2011; Möckel et al., 2012). A recent cryo-EM structure of *E. coli* MR (called SbcCD) bound to a double-stranded DNA (dsDNA) substrate revealed a structure where the ADP-bound Rad50s are associated and the Mre11 dimer has moved to one side to interact asymmetrically with the two Rad50s and the dsDNA substrate (Käshammer et al., 2019). In addition, we have previously used Luminescence Resonance Energy Transfer (LRET) experiments to illuminate the presence of a ‘partially open’ conformation in a truncated construct of hyperthermophilic *P. furiosus* MR (Pf MR^NBD^) in both ATP and ATP-free conditions (Boswell et al., 2020). Thus, a variety of functionally relevant structures of the MR complex may exist in solution. As LRET has been successfully used to characterize the interactions of NBDs in several ABC ATPase membrane proteins (Cooper and Altenberg, 2013; Zoghbi et al., 2017, 2012; Zoghbi and Altenberg, 2018), we significantly extended our initial LRET studies on the Pf MR^NBD^ complex to further characterize this partially open conformation. Multiple LRET probes were introduced throughout the Rad50 NBD to determine a network of distances between residues across the Rad50-Rad50 interface (Fig. 1B). These LRET-determined distances were then used as unambiguous distance restraints in the molecular docking program HADDOCK (Dominguez et al., 2003; Van Zundert et al., 2016) to obtain models of the ATP-bound MR^NBD^ complex. Here, we present structural models of three distinct conformations of the ATP-bound Pf MR^NBD^ complex: closed, partially open, and open. LRET experiments on full-length Pf MR, where Rad50 contains the coiled-coil domains and an apical zinc hook dimerization motif, confirmed that these conformations are also observed for the complete MR complex. Site-directed mutagenesis was used to disrupt specific conformations, and the effects on Mre11 and Rad50 activities validated our models and put them into a functional context. Small-angle X-ray scattering (SAXS) was also employed to confirm these three conformations of ATP-bound MR^NBD^ in solution and to assign approximate populations to each. In conclusion, we have combined orthogonal biophysical and computational methods to describe three distinct global conformations of ATP-bound Pf MR in solution and demonstrate that they offer valuable insight into how MR functions along the DNA.

## Results

### Multiple LRET probe positions provide a network of measurements

LRET experiments employ a luminescent lanthanide donor and fluorophore acceptor pair with an appropriate Förster radius (R_0_)(Zoghbi and Altenberg, 2018). LRET probes are introduced into a protein most easily through a thiol-maleimide reaction with a unique cysteine. For Pf Rad50^NBD^, where the coiled-coil domains are truncated, single cysteines were introduced into the naturally cysteine-less construct, and MR^NBD^ activity was tested to ensure an active complex (Supplemental Fig. S1). Mutations were made primarily in loop regions to minimize disruptions to the protein fold. In all, six separate single cysteine mutations were made throughout Rad50^NBD^ (Fig. 1B). These cysteines were subsequently labeled with thiol-reactive LRET donor or acceptor molecules. To make the MR^NBD^ complex for LRET experiments, equimolar amounts of donor-labeled (Tb^3+^-chelate) Rad50^NBD^ and acceptor-labeled (Bodipy FL or Cy3) Rad50^NBD^ were mixed with twice the molar ratio of Mre11 so that 50% of the resulting M_2_R_2_ complexes had one donor and one acceptor fluorophore (on separate Rad50 protomers). Not only were identical cysteines mixed within a complex (e.g., Tb^3+^-S13C and Bodipy-S13C), but complexes were also made where cysteine mutants were mixed with other cysteine mutants (e.g., Tb^3+^-S13C and Bodipy-L51C). Additional distance measurements were obtained for a given LRET pair by changing the identity of the acceptor, as the Förster radius for Tb^3+^ and Cy3 (61.2 Å) is longer than that of Tb^3+^ and Bodipy FL (44.9 Å). Finally, donor-and acceptor-labeled cysteines within mixed pairs were swapped (e.g., Tb^3+^-S13C and Bodipy-L51C versus Tb^3+^-L51C and Bodipy-S13C) for added confidence in measurements. In total, 20 different cysteine pairs resulted in 50 unique samples that gave 54 total measured distances between the two Rad50 protomers in the MR^NBD^ complex (Supplemental Table S1).

### LRET measurements reveal three distinct sets of distances

Following laser excitation of the Tb^3+^- chelate moiety and a 200 μsec delay, donor-sensitized Bodipy FL or Cy3 fluorescence emission decay curves were collected for each of the MR^NBD^ LRET samples at 50 °C in the presence of 5 mM Mg^2+^ and 2 mM ATP (Fig. 1C). Under these conditions, Rad50^NBD^ should be >99% bound to ATP as the K_D_ for ATP is ∼3 μM and there is no measurable ATP hydrolysis in 1 h at 50 °C. In multi-exponential fits, the emission decays were best described by two or three exponentials depending on the identity of the LRET pair (see Methods). In all cases the first lifetime (<100 μs) is a function of instrument response time and was discarded (Cooper and Altenberg, 2013; Zoghbi et al., 2017, 2012). The Tb^3+^-chelate luminescence decays were also recorded at each probe position in donor-only labeled MR^NBD^ complexes. As expected, the value of the Tb^3+^-chelate lifetime changed with the local environment of each cysteine. Using these Tb^3+^-chelate donor lifetimes, combined with the donor-sensitized acceptor lifetimes and the R_0_ of the dye pair in the sample, distances were calculated between probes for each LRET pair.

For the majority of the 20 cysteine pairs analyzed, combining the data for the Bodipy/Tb^3+^- and Cy3/Tb^3+^-labeled samples gave three distinct distances. For 10 of the pairs (e.g., L51C-L51C), the longer distance (d2^Bodipy^) in the Bodipy/Tb^3+^ samples matched the shorter distance (d1^Cy3^) in the Cy3/Tb^3+^ samples, and the Cy3/Tb^3+^ samples gave a second, longer distance (d2^Cy3^) (Fig. 1D and Supplemental Table S1). This longer distance became ‘visible’ in samples where Cy3 was the acceptor because the R_0_ for Tb^3+^ and Cy3 is longer. For four of the pairs (e.g., L51C-A66C), the one Bodipy/Tb^3+^ distance did not overlap with the two Cy3/Tb^3+^-determined distances, and the combined data resulted in three distances. For two pairs (A66C-A66C and S93C-S93C), only one distance was seen in the Bodipy/Tb^3+^ data, while the Cy3/Tb^3+^ data contained two distances. In these samples, the d^Bodipy^ matched d1^Cy3^ for a total of two distances. And finally, for four pairs (e.g., S13C-S93C) only Cy3/Tb^3+^ samples were made resulting in two distances. Together, these data illuminate the presence of a third solution state in addition to the closed and partially open.

To confirm that the distances observed in MR^NBD^ were the same in full-length MR, all of the cysteine mutations were also introduced into full-length Pf Rad50. We previously reported that the two native cysteines in the zinc hook motif of full-length Rad50 are not efficiently labeled by the LRET fluorophores and do not result in LRET donor-sensitized acceptor signal (Boswell et al., 2020). Tb^3+^- chelate donor only lifetimes measured in these mutants were identical to those measured for the Rad50^NBD^ construct, indicating that the local environments of the introduced cysteines do not change between full-length and NBD constructs. Unfortunately, because full-length Rad50 dimerizes at the zinc hook, labeled cysteine mutants could not be mixed and only “identity” LRET pairs (e.g., Bodipy-L51C and Tb^3+^-L51C) could be made. Nonetheless, for all LRET probe positions measured, the distances between full-length MR cysteine pairs were within a few Ångströms of those measured in MR^NBD^ (Fig. 1D).

### HADDOCK models of three MR^NBD^ conformations

The measured LRET distances were input as unambiguous restraints in the HADDOCK molecular docking program(Dominguez et al., 2003; Van Zundert et al., 2016), defining the Cβ-Cβ distance between the LRET-labeled residues. The unambiguous restraints included symmetrical distances for each LRET pair (e.g., both Rad50a protomer L51 to Rad50b protomer S13 and Rad50a S13 to Rad50b L51). These unambiguous restraints were used to dock two Rad50^NBD^ protomers with one Mre11 dimer in a three-body docking simulation.

The ‘closed’ HADDOCK model fit very closely to the measured LRET distances (Fig. 2A, Supplemental Movie S1, and Supplemental Table S2). HADDOCK returned 190 structures in five clusters, with 164 in the top-scored cluster. The Rad50 dimer formed in this model is nearly identical to the AMPPNP-bound dimer structure of Pf Rad50^NBD^ (PDB: 3QKU, all-atom root-mean-square deviation [RMSD] = 1.11 Å) (Williams et al., 2011). Except in two cases, the Cβ-Cβ distances between the Rad50 LRET pairs in the HADDOCK model were within ±5 Å of their respective unambiguous LRET restraint. LRET pairs A66-A66 and A66-S93 deviated more significantly with differences of 7.4 and 11.0 Å, respectively. This deviation could arise from slight differences in the loop structures between the solution (LRET) and crystal states (i.e., input PDB; note that HADDOCK does not move backbone atom positions during model refinement). Specifically, we observed the expected relative positions within the associated Rad50s for residues in the Walker A motif (N32) of one protomer and the D-loop (D829) and signature motif (S793) of the other (Fig. 2D, top and movie S1). In this closed model, Rad50 interacts with Mre11 via the capping domain and along the top of the nuclease domain as observed in the *T. maritima* (PDB: 3THO) (Möckel et al., 2012), *M. jannaschii* (PDB: 3AV0) (Lim et al., 2011), and *E. coli* (PDB: 6S6V) (Käshammer et al., 2019) nucleotide-bound structures (with Cα RMSDs of 4.9 Å, 1.9 Å, and 4.7 Å, respectively) (Supplemental Fig. S2A). Like the *M. jannaschii* structure, each Rad50 protomer makes contact with only one of the Mre11 capping domains mainly through interactions between capping domain β18 and Rad50 Lobe II αE and β8-10. In total, eight and four ionic or hydrogen bond interactions are made between Rad50 and the Mre11 capping and nuclease domains, respectively. In particular, unique contacts not seen in the other two conformations are made between Rad50 E831 and Mre11 H17 in the nuclease domain and Rad50 E758/E761 and Mre11 Y325/K327 in the capping domain. The combination of these interactions occludes the Mre11 nuclease active site for dsDNA, as previously described (Lim et al., 2011; Möckel et al., 2012).

**Fig. 2.**
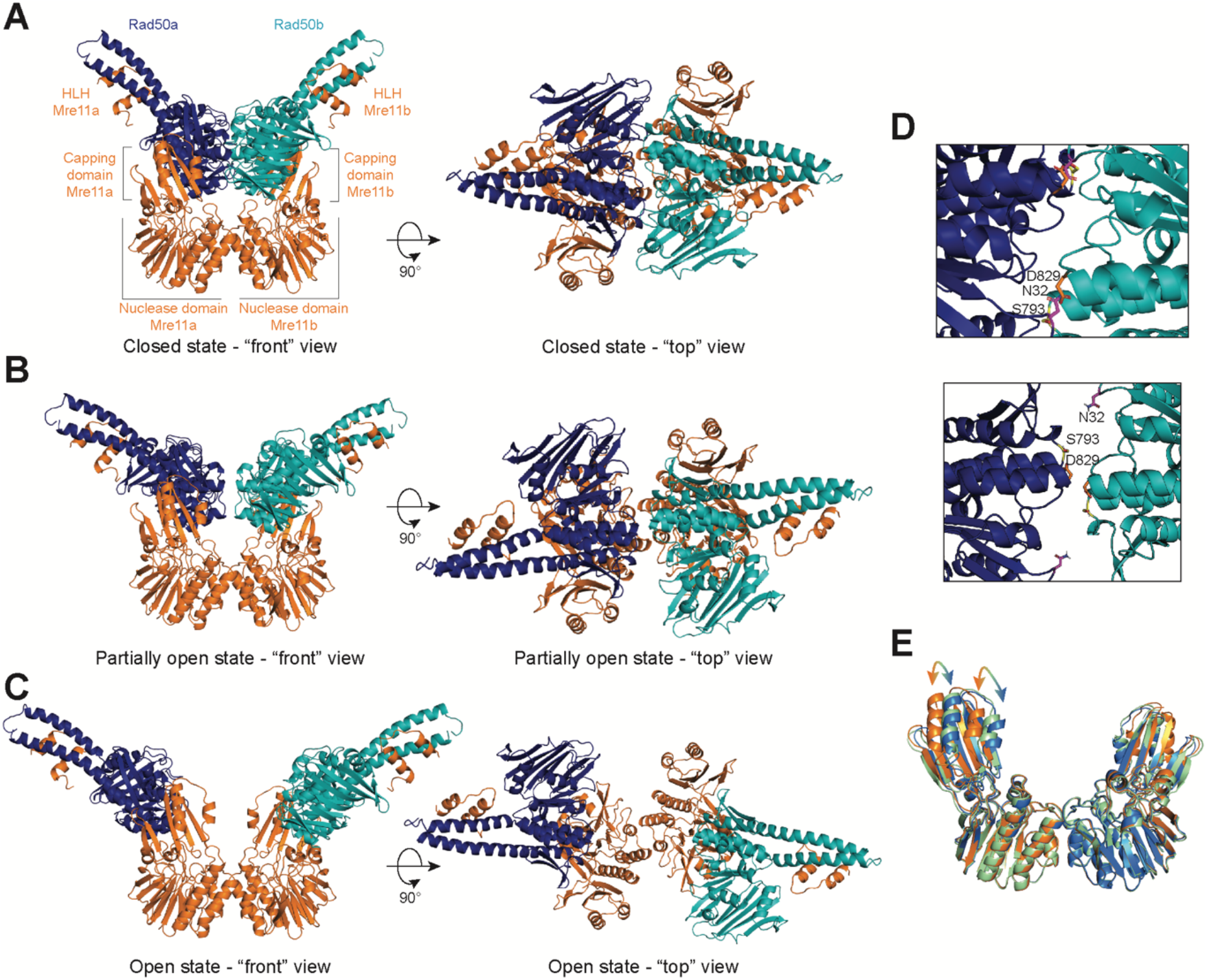
The Pf MR^NBD^ ATP-bound complex has at least three conformations in solution. HADDOCK structural models of the (A) closed, (B) partially open, and (C) open MR^NBD^ complex. (D) The Rad50-Rad50 interface with the Walker A residue (N32, magenta) from Rad50b and the Signature helix (S793, yellow) and D-loop (D829, orange) residues of Rad50a indicated in the closed (top) and partially open (bottom) conformations. (E) Overlay of the Mre11 dimers from the closed (orange), partially open (green), and open (blue) HADDOCK models showing that the capping domain moves out (arrows) to accommodate the associated Rad50s in the closed conformation.

The ‘partially open’ HADDOCK simulation returned nine clusters of models, and analysis of the top four clusters concluded that the distances between LRET pairs for each cluster had an average standard deviation of ∼0.3 Å when compared among the top three clusters and increased to ∼0.7 Å when adding the fourth (Fig. 2B, Supplemental Movie S1, and Supplemental Table S2). 13 out of 20 of the Cβ-Cβ distances in the HADDOCK model are within ±5.7 Å of the measured LRET distances (Supplemental Table S2). Interestingly, all of the pairs with larger (>5.7 Å) deviations included either A66 or S93, again suggesting that the position of those loops differ between the crystal and solution conditions. In the partially open model, only 4.2 Å separates the Rad50 protomers at their closest point between the two D829 residues (Cα-Cα distance). In this conformer, the Walker A/D-loop interactions are no longer formed between the Rad50 protomers, as the Walker A motifs have rotated out of the Rad50-Rad50 interface (Fig. 2D, bottom and Supplemental Movie S1). Instead, the D-loop and signature motifs face one another and are stacked in the interface. Although there are still significant interactions between the Rad50 protomers and the Mre11 capping and nuclease domains, several of which are maintained from the closed conformation, the partially open conformation has rotated by 22° to interact differently with the capping domain compared to the closed structure (Supplemental Fig. S2B). Specifically, interactions with the W308-D313 loop in the Mre11 capping domain are similar between the two conformations, but the Rad50 interactions with β18 have been broken in partially open, and Rad50 is now interacting with Mre11 β16 and β17 residues instead. Moreover, the movement of Rad50 has allowed the Mre11 capping domains to rotate inward toward the nuclease domains (Fig. 2E). We observe a total of nine ionic or hydrogen bond interactions between the Mre11 capping domain and Rad50 and five between the nuclease domain and Rad50. In the capping domain, unique interactions occur between Mre11 K277 and Rad50 E750 and Mre11 R303 and Rad50 E754. Even with these contacts between Rad50 and Mre11, DNA could access the nuclease active site of Mre11.

Finally, HADDOCK returned 14 clusters of models for the ‘open’ complex, and analysis of the top four clusters showed that the distances between LRET pairs for each cluster had an average standard deviation of ∼0.7 Å among the top three clusters and ∼1.2 Å when adding the fourth. Seven out of 10 of the Cβ-Cβ distances in the top cluster are within ±6 Å of the input LRET unambiguous restraints whereas the remaining three are within ±9.3 Å (Fig. 2C, Supplemental Movie S1, and Supplemental Table S2). The Rad50 protomers have moved apart considerably (∼41 Å D829-D829 Cα-Cα distance). Moreover, the orientation of the Rad50 protomers with respect to Mre11 has changed significantly (Supplemental Fig. S2C) having rotated 13° and the base of the coiled-coil translated ∼23 Å away from the partially open model. Because of this rotation and translation, there are minimal contacts (only two) with the Mre11 capping domain, which now occur between Rad50 β10 and Mre11 β16 and the C-terminus of this construct. Now, six ionic interactions are formed between the Mre11 nuclease domain and Rad50. Within the nuclease domain, unique interactions occur between Mre11 R177 and E181 on helix αE with Rad50 E841 and R842 and between Mre11 E152 on helix αD and Rad50 K860. In the capping domain, Mre11 K279 makes a unique interaction with Rad50 E783. With Rad50 rotated fully away, the capping domains move even closer to the nuclease domain and both Mre11 nuclease active sites are now fully accessible to dsDNA for exonuclease activity.

To ensure that the unambiguous distance restraints obtained from one probe position were not dominating the structure calculations, HADDOCK runs were performed with systematic dropouts of all restraints calculated from a specific cysteine position. For example, for the L51C probe position, L51C-S13C, L51C-L51C, L51C-A66C, L51C-S93C, L51C-N774C, and L51C-V866C distances were all removed from the unambiguous restraints, and HADDOCK runs were repeated for each of the three sets of distances (closed, partially open, and open). In general, none of the dropouts appreciably changed the overall conformation of any of the states (Supplemental Fig. S3).

### Destabilizing solution state conformers alters MR activity

Next, to decrease the stability of one or two conformations over the others, charge reversal mutations were made to several Mre11 residues that directly interact with Rad50. As there are a handful of shared interactions between Mre11 and Rad50 in the various conformation combinations, it was impossible to completely disrupt a given state. These Mre11 mutants were combined with full-length Rad50 to make MR complex, and then both Mre11 and Rad50 activities were tested (Fig. 3). To assay Mn^2+^-dependent Mre11 3’-to-5’ exonuclease activity, two dsDNA substrates were interrogated: Exo2 and Exo11, which have a fluorescent 2-aminopurine incorporated as the second or eleventh nucleotide from the 3’-end of one strand, respectively. Although MR does not require ATP to cleave the Exo2 substrate, ATP is required for Exo11 as the enzyme needs to progress eleven base pairs along the DNA duplex to reach the position of the fluorescent nucleotide (Boswell et al., 2020; Herdendorf et al., 2011).

**Fig. 3.**
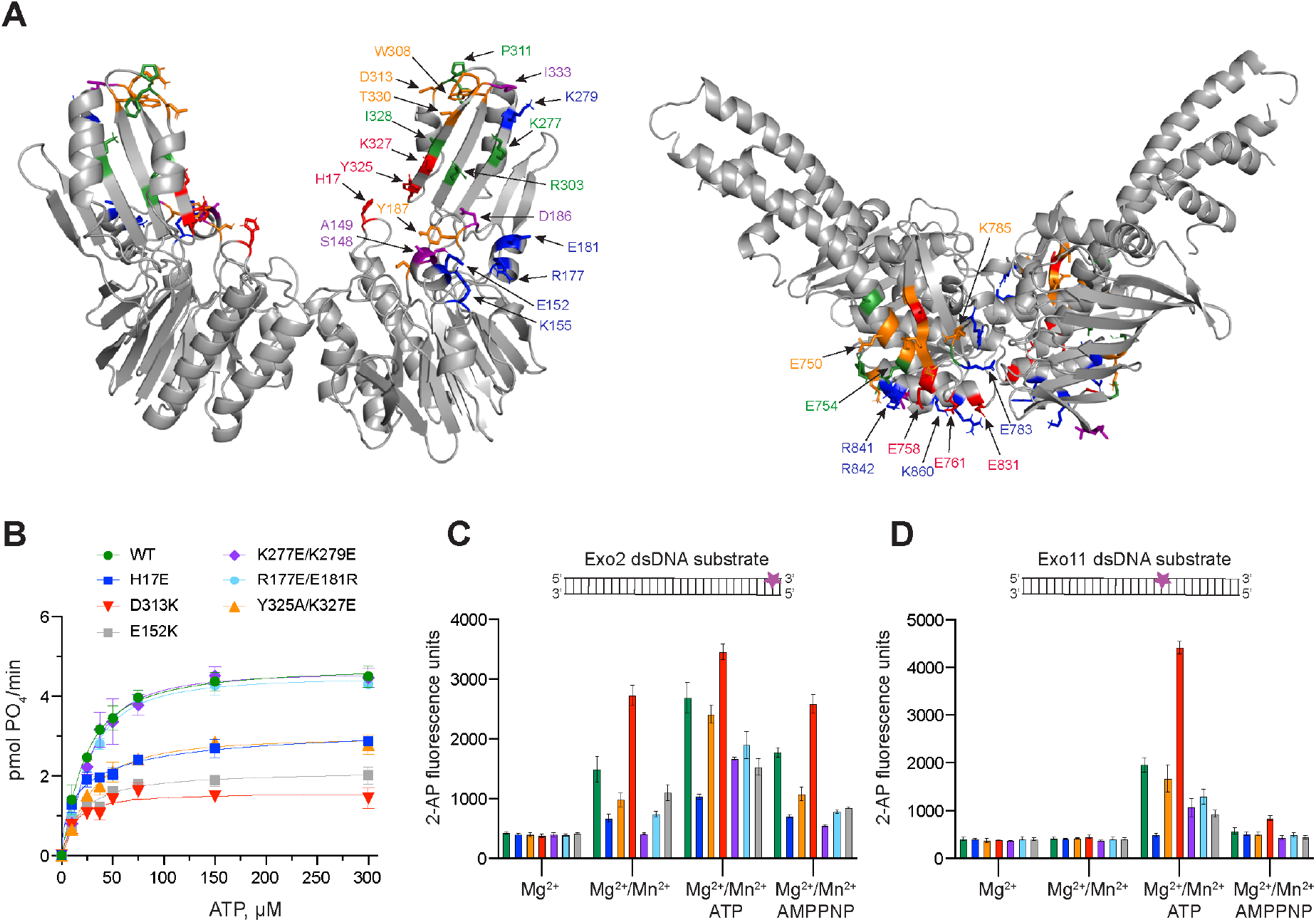
Partially open and open conformations of the MR complex are important for exonuclease activity. (A) Mre11 (left) and Rad50 (right) dimers showing residues involved in protein-protein interactions only in closed (red), only in partially open (green), and only in open (blue) conformations or common to closed and partially open (orange) or common to all three (purple). (B) ATP hydrolysis for MR complexes containing the indicated Mre11 mutants. (C) and (D) MR complex exonuclease activity on the Exo2 (C) or Exo11 (D) dsDNA substrates. Position of fluorescent 2-AP is indicated with a star on the cartoon of each substrate. Mre11 mutant colors are same as in (B). Data are the mean and standard deviation of n ≥3 replicates.

In the closed conformation, Mre11 nuclease domain residue H17 interacts with Rad50 E831 (Supplemental Fig. S4A). MR H17E had ∼30% of wild type exonuclease activity on Exo2 DNA but does not cleave Exo11 DNA. Williams et al. identified H17 as a ‘wedge residue’ that helps to unwind the dsDNA helix (Williams et al., 2008). The observation that processive MR exonuclease activity is hindered in the H17E mutant supports that function. MR H17E decreased the V_max_ of ATP hydrolysis to ∼60% of wild type MR, indicating that this residue also assists in stabilizing the closed conformation from which hydrolysis proceeds. Mre11 Y325 and K327 are located in the capping domain on β18 and interact with E761 and E758 of Rad50, respectively, in the closed conformation (Supplemental Fig. S4A). In the presence of ATP, the Y325A/K327E double mutant had ∼85% of the exonuclease activity of wild type MR, but only ∼50% of the activity for the Exo2 substrate without ATP. MR Y325A/K327E decreased the V_max_ of ATP hydrolysis to the same level as MR H17E. Therefore, destabilizing the closed conformation through the Mre11 H17E and Y325A/K327E mutants had the predictable effect of decreasing Rad50 ATP hydrolysis.

Mre11 D313 contacts Rad50 K785 in both the closed and partially open conformations (Supplemental Fig. S4A and S4B). D313 and its neighboring residues in a loop at the top of the capping domain might be acting as a pivot point for Rad50 to rotate between the two states. Surprisingly, MR D313K had significantly increased exonuclease activity (Fig. 3). In fact, a ∼2-fold increase in activity was observed for Exo2 in the absence of ATP; thus, we hypothesize that destabilizing both the closed and partially open conformations increases the population of the open conformation that accommodates dsDNA substrate. Activity against the Exo11 substrate increased ∼2.5-fold in this mutant. In contrast to its high exonuclease activity, MR D313K reduced the V_max_ for ATP hydrolysis by more than 50%, which was expected since the closed state is destabilized.

In the partially open complex, Mre11 K277 in β16 interacts with Rad50 E750, which is in αE at the base of a coiled-coil, whereas in the open complex, Mre11 K279 also in β16 interacts with Rad50 E783 in the short loop between β9 and β10 (Supplemental Fig. S4B and S4C). As these two residues are close in sequence space, a double mutant was constructed to destabilize both partially open and open conformations. The most striking feature of MR K277E/K279E was that it had no nuclease activity on Exo2 without ATP but increased to ∼50% of wild type levels when ATP was added (Fig. 3). No other mutant tested required ATP for the Exo2 substrate, implying that it requires hydrolysis to open the complex before dsDNA can bind. Moreover, this mutant displayed the lowest activity in the presence of AMPPNP suggesting that the combination of the mutations and non-hydrolyzable analog effectively stabilized the closed state, fully occluding the dsDNA substrate. On the Exo11 substrate, the activity was also ∼50% of wild type. This double mutant has no effect on ATP hydrolysis activity, since the closed conformation can readily form.

Finally, mutants were made for Mre11 R177, E181, and E152, which all make unique contacts in the open conformation (Supplemental Fig. S4C). These residues are along the “top edge” of the Mre11 nuclease domain in αD and αE, and R177/E181 and E152 are structurally homologous to part of the so-called “latching loop” and “fastener,” respectively, of the *E. coli* SbcCD cryo-EM “cutting state” structure (Käshammer et al., 2019). The MR R177E/E181R double mutant had only ∼60% of wild type exonuclease activity but wild type ATP hydrolysis activity, which is consistent with there being less open and more closed conformation. On the other hand, MR E152K, which interacts with Rad50 K860 in αE, showed more impaired exonuclease activity than R177/E181, but surprisingly decreased the V_max_ for ATP hydrolysis by more than 50% (Fig. 3). As this mutant should readily form the closed conformation, this result was unexpected. Like the K227E/K279E mutant, both MR R177E/E181R and MR E152K had very little Exo2 activity in the presence of AMPPNP, again suggesting a complete stabilization of the closed conformation.

### SAXS corroborates HADDOCK MR^NBD^ models

To further validate the multiple conformations indicated by the LRET data, SAXS data was collected for samples of MR^NBD^ with and without ATP. The SAXS profiles for wild type MR^NBD^ were compared to our three models using the *FoXS* webserver (Schneidman-Duhovny et al., 2016, 2013), which calculates the SAXS profile from an input structural model and compares that to an experimental SAXS profile. The ATP-bound MR^NBD^ experimental SAXS profile fit best to the partially open model (χ^2^ = 0.34) followed by the closed model (χ^2^ = 0.51), based on the single-state χ^2^ values (Fig. 4). The open model did not fit well to the experimental SAXS data (χ^2^ = 13.6). Conversely, for the ATP-free MR^NBD^ SAXS profile, a reasonable fit was only obtained with the open model (χ^2^ = 1.26). Next, *MultiFoXS* was used to fit the ATP-bound MR^NBD^ SAXS profile to either two or three of the LRET-derived models simultaneously. The fit to two models improved the χ^2^ significantly and combined populations of open (19%) and closed (81%) conformations (Fig. 4B). The fit using all three models did not improve the χ^2^ but did assign populations of 15% open, 18% partially open, and 67% closed. The two-state fit to the apo MR^NBD^ SAXS profile only slightly improved the χ^2^ (1.26 *vs* 1.05) and gave 90% open and 10% closed, while the 3-state fit gave 89% open, 5% partially open, and 6% closed again without improving χ^2^ further. Thus, the SAXS data supports the three states observed in LRET data and demonstrates the expected shift in population to the closed form in the presence of ATP.

**Fig. 4.**
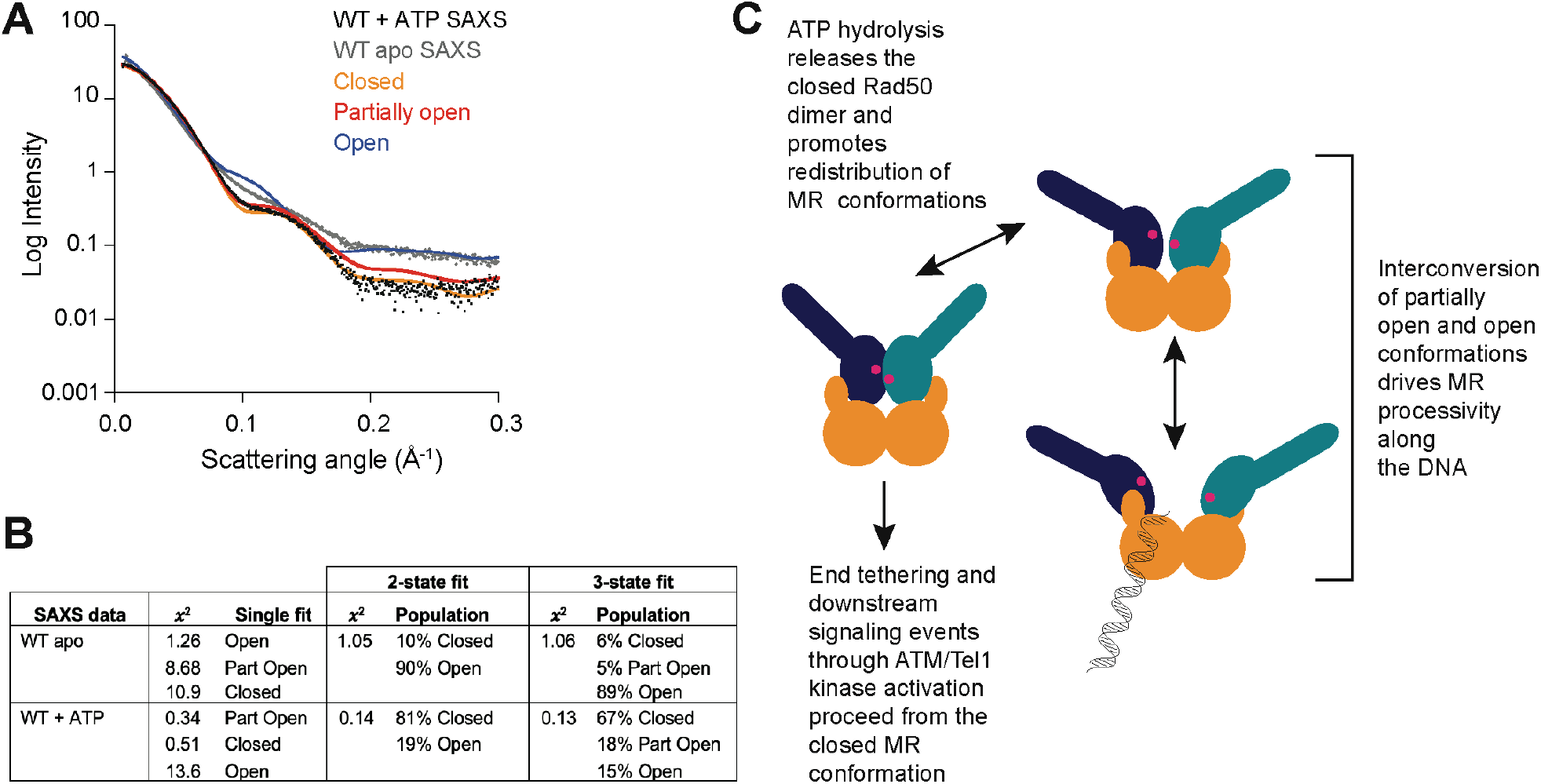
All three MR conformations play a role in the function of the complex. (A) Experimental SAXS curves for apo (grey) and ATP-bound (black) MR^NBD^ superimposed with FoXS-calculated theoretical SAXS curves for the closed (orange), partially open (red), and open (blue) HADDOCK models. (B) Goodness of fits of experimental to theoretical SAXS curves calculated by FoXS and MultiFoXS. For fits to two or three of the models, the populations from the fit are shown. (C) Proposed model of the functional role of conformational heterogeneity in the MR complex.

## Discussion

The disulfide bonds and non-hydrolyzable ATP analogs used in all of the closed MR^NBD^ crystal structures and the selection for closed particles during the *E. coli* SbcCD cryo-EM model refinement imply that there is more than one conformation of ATP-bound MR. Nevertheless, the ATP-bound closed form has been referred to as the “resting” state given the cellular concentration of ATP and the K_D_ for ATP-binding to Rad50 (Käshammer et al., 2019). Even so, both closed and partially open conformations were also identified in apo MR^NBD^ (Boswell et al., 2020). In fact, the existing structural data (Deshpande et al., 2014; Williams et al., 2011) and the LRET-derived HADDOCK models presented here provide evidence for at least three conformations for Pf MR^NBD^ in solution: closed, partially open, and open. As LRET is a distance-dependent phenomenon and distances could not be measured for probes more than ∼85 Å apart, we did not obtain data for the extended conformation. Likewise, there was no evidence of MR^NBD^ complexes containing dissociated Mre11 dimers (Käshammer et al., 2019; Saathoff et al., 2018), since the LRET probes on the Rad50 NBDs would also be too far apart in that scenario. Interestingly, this is not the first time that conformations with these names have been suggested for MR. Williams and co-workers(Williams et al., 2011) found that a combination of molecular dynamics (MD)-simulated conformational models best described their ATP-free and ATP-bound Pf MR^NBD^ SAXS curves. Although those MD models do not resemble the models presented here, they nevertheless opened the door for conformational heterogeneity in MR, a possibility also proposed by others (Deshpande et al., 2014; Lafrance-Vanasse et al., 2015).

Our characterization of Mre11-Rad50 interaction mutants in full-length MR complexes, along with the 41 Å separation between Rad50 monomers, suggests that only the open conformation can accommodate dsDNA in a productive orientation. When the stability of both the open and partially open conformations are compromised (K277E/K279E), MR cannot cleave the Exo2 substrate without ATP, which once hydrolyzed likely results in destabilization of the closed conformation for enough time to allow dsDNA to bind. The R177E/E181R mutant only disrupts the open conformation, but unlike K277E/K279E, there is some residual Exo2 activity without ATP. Perhaps this is because the partially open conformation can form and occasionally flexes open enough for dsDNA to bind. Notably, neither of these mutants that should form more of the closed conformation increase ATP hydrolysis, since that reaction is highly regulated by allostery within Rad50 (Boswell et al., 2018; Deshpande et al., 2014; Williams et al., 2011). On the other hand, D313K, which destabilizes both the closed and partially open conformations to form the open conformation that binds dsDNA, has the highest levels of exonuclease activity, cleaving the Exo2 substrate readily without ATP. Additionally, this mutant is able to process along the DNA even though it has significantly decreased ATP hydrolysis, suggesting that although hydrolysis is required to release the closed state and redistribute the population of the conformations, it may not be specifically required for exonuclease activity. This hypothesis is further supported when looking at all of the Exo2 reactions without ATP (i.e., Mg^2+^/Mn^2+^): with the exception of the mutant that has a stabilized closed form (K277E/K279E) all of the MR complexes can cleave DNA. Furthermore, the D313K mutant is in a loop that may act as a pivot between the closed and partially open conformations. By hampering the ability of MR to pivot here, there may be more interconversion between partially open and open conformations, which, we hypothesize, drives MR processivity along the DNA.

In order for the Mre11 dimer to accommodate the Rad50 dimer in the closed conformation, the capping domain has flexed outward ∼5 Å (Fig. 2E and Supplemental Movie S1) from its position in the partially open and open conformations. Additional capping domain motions are also observed when transitioning between partially open and open conformers. This movement within the capping domain is reminiscent of motions previously observed in structures of the Pf Mre11 dimer bound to DNA substrates that mimicked either a DNA DSB or a stalled replication fork (Williams et al., 2008). Thus, our MR models further confirm that motions within the Mre11 capping domain coupled with the H17 wedge residue are important in DNA unwinding and processive exonuclease activity.

In summary, our data suggest a model for MR activity where the ATP-bound closed, partially open, and open conformations exist in equilibrium (Fig. 4C). Initial recognition of the DNA DSB occurs with the open state, where at least the first two nucleotides can be excised. Resection proceeds as MR cycles between the open and partially open conformations. Since the D313K mutant can perform processive exonuclease activity with very low ATP hydrolysis activity, we suggest that the closed form may not be necessary for exonuclease function per se. Instead, the closed form could serve to reset the equilibrium of the three states once bound ATP is hydrolyzed and products are released. Importantly, the closed conformation must form in order for MR to be functional, as it is required for downstream signaling through ATM. Our model is supported by several recently reported cancer-associated MR DNA DSB separation-of-function mutants which maintain Mre11 nuclease function while losing the ability to signal the presence of the DNA DSB through ATM or vice versa (Al-Ahmadie et al., 2014; Chansel-Da Cruz et al., 2020; Hohl et al., 2020). Thus, the conformational heterogeneity we describe here has important functional consequences for DNA double strand break repair.

## Methods

### Plasmids, protein expression, and purification

Plasmids for *P. furiosus* full-length Rad50, Rad50 nucleotide binding domain (Rad50^NBD^: aa1-195; GGAGGAGG linker; aa709-882), and Mre11 protein expression in *E. coli* were previously described (Boswell et al., 2020, 2018). Point mutations were introduced using a modified Quikchange (Strategene) approach and were verified by Sanger sequencing. Protein expression and purification were performed as previously described for full-length Rad50 (Boswell et al., 2020), Rad50^NBD^ (Boswell et al., 2018), and Mre11 (Rahman et al., 2020).

### ATP hydrolysis assay

Rad50 ATP hydrolysis assays were performed essentially as described by Boswell *et al*. (Boswell et al., 2018). 0-300 μM ATP was titrated into microfuge tubes containing either 2.5 μM MR^NBD^ complex or 2 μM MR complex and 50 mM Tris, 80 mM NaCl, 1% glycerol, 5 mM MgCl_2_, pH 7. Reactions without protein were included for each ATP concentration to control for ATP degradation and PO_4_ contamination. 60 μL reactions were incubated at 65 °C for 60 min after which the tubes were placed on ice. 50 μL of each reaction was then transferred to the wells of clear, flat-bottom 96-well plates and 100 μL of cold BIOMOL Green (Enzo Lifesciences) colorimetric reagent was added. After a 30 min incubation at room temperature to allow the color to develop, the amount of inorganic phosphate released by hydrolysis was quantified using the absorbance mode on a Synergy Neo2 multi-mode plate reader. BIOMOL Green signal (A_640_) was corrected by subtracting the A_640_ values of the ATP only reactions at each ATP concentration and then transformed into pmols of PO_4_ released/min based on a PO_4_ standard curve incubated in BIOMOL Green reagent for 30 min at room temperature. Plots of PO_4_ released/min (v_0_) versus ATP concentration were fit to the Michaelis-Menten equation including a Hill coefficient (n).

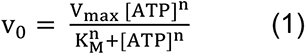

### Exonuclease assays

Exonuclease activity was assayed by monitoring the fluorescence of a 2-aminopurine (2-AP) nucleotide incorporated into a dsDNA substrate. 2-AP fluorescence is quenched within the base pair, and release of the 2-AP from the duplex by Mre11 exonuclease activity results in an increase in the 2-AP fluorescence signal. The Exo2 substrate was formed by annealing equimolar amounts of 5’-GGCGTGCCTTGGGCGCGCTGCGGGCGG[2-AP]G-3’ and 5’- CTCCGCCCGCAGCGCGCCCAAGGCACGCC-3’ DNA oligos. For LRET cysteine mutants, 30 μL exonuclease reactions were mixed in microfuge tubes and contained 0.5 μM MR^NBD^ complex and 1 μM Exo2 dsDNA substrate in 50 mM Hepes, 100 mM NaCl, 5 mM MgCl_2_, 0.1 mM EDTA, 1% glycerol, 1 mM TCEP, pH 7. 1 mM MnCl_2_ was added to the reactions as indicated. After a 45 min incubation at 60 °C, the tubes were removed from the heat block and spun. 25 μL of each reaction was transferred to black, flat-bottom 384-well plates and 2-AP fluorescence was quantified by the Synergy Neo2 plate reader (ex310/em375). For the Mre11-Rad50 interaction mutants, exonuclease activity was assessed for MR complexes made with full-length Rad50. Reactions with 0.5 μM MR complex and 1 μM Exo2 were assembled as above, except 1 mM MnCl_2_, 1 mM MnCl_2_/1 mM ATP, or 1 mM MnCl_2_/1 mM AMPPNP were added as indicated and reactions were incubated at 60 °C for 15 min. In addition to the Exo2 substrate, a second substrate, Exo11, was also employed for MR: 5’-GGCGTGCCTTGGGCGCGC[2- AP]GCGGGCGGAG-3’ annealed to 5’-CTCCGCCCGCTGCGCGCCCAAGGCACGCC-3’. Exo11 reactions were incubated at 60 °C for 30 min.

### Labeling Rad50 cysteine mutants with LRET probes

Thiol-reactive Tb^3+^ chelate DTPA-cs124-EMPH (Lanthascreen, Life Technologies, Inc.) was used as the LRET donor for all samples and Bodipy FL maleimide (Invitrogen) or Cyanine3 (Cy3) maleimide (GE Healthcare) were used as acceptor fluorophores. 50 μM purified Rad50^NBD^ containing a single cysteine was combined with a 2-fold excess of one of the labels in degassed Labeling Buffer (25 mM Tris pH 8, 100 mM NaCl, 10% glycerol, 1 mM TCEP) in 100 μL reactions. The reaction was incubated in the dark at room temperature for 1.5 to 2 h. Labeling of full-length Rad50 cysteine mutants followed the same protocol, except the donor and one acceptor label were added simultaneously to each reaction (2-fold molar ratio of each). Unreacted label was removed by running the reaction over a Superdex 200 Increase 10/300 GL column (GE Healthcare) equilibrated with 25 mM Hepes, 200 mM NaCl, 10% glycerol, 1 mM TCEP, pH 7. Eluted protein was concentrated using centrifugal concentrators (VivaSpin, Sartorius) and then assessed for successful labeling by measuring fluorescence using the following settings and the monochromator in the BioTek Synergy Neo2 plate reader: ex337/em490 for DPTA-Tb^3+^, ex485/em515 for Bodipy FL, and ex535/em570 for Cy3. Concentrated labeled proteins were aliquoted, frozen in liquid nitrogen, and stored at −80 °C.

### LRET data collection and analysis

For LRET experiments, 40 μL mixtures of 1 μM Mre11 + 0.5 μM Tb^3+^-labeled Rad50^NBD^ + 0.5 μM Bodipy- or Cy3-labeled Rad50^NBD^ were heated at 60 °C for 15 min to form the MR^NBD^ complex. Full-length MR LRET mixtures were 1 μM Mre11 + 1 μM dual-labeled (donor + acceptor) Rad50. Each LRET sample was subsequently diluted to 160 μL (0.25 μM complex) with LRET Buffer (50 mM Hepes, 100 mM NaCl, 5 mM MgCl_2_, 0.1 mM EDTA, 1% glycerol, 1 mM TCEP, pH 7) before 2 mM ATP was added. MR^NBD^+ATP and MR+ATP LRET samples were heated again for 15 min at 50 °C before 150 μL were transferred into a 3-mm pathlength cuvette. All LRET data were collected at 50 °C. Donor and acceptor lifetimes were calculated from intensity decays measured with an Optical Building Blocks phosphorescence lifetime photometer (EasyLife L). LRET samples were excited through a narrow band 335 nm filter (Semrock FF01-335/7) and after a 200 μs delay (to allow the decay of sample autofluorescence, emission due to direct excitation of the acceptor, and scattering of the excitation pulse), donor emission intensity was collected for Tb^3+^ at 50 Hz through a 490/10 nm band-pass filter (Omega Optical) and donor-sensitized acceptor emission intensities were collected at 100 Hz through 520/10 nm (Bodipy FL) or 570/10 nm (Cy3) band-pass filters. PTI FeliX32 software was used to fit the Bodipy FL and Cy3 emission decay curves to either a two- or three-exponential function depending on the identity of the LRET pair. For samples where the distance between the LRET pair in the nucleotide-bound crystal structure was ≤42 Å, the data fit well to three exponentials for both Bodipy- and Cy3-labeled samples. For samples where the LRET pair was separated by more than 45 Å, the Bodipy emission decays fit to only two exponentials while the Cy3 decays fit to three. Lifetime distributions shorter than ∼100 μs were discarded as these are largely a function of the instrument response time (Cooper and Altenberg, 2013; Zoghbi et al., 2017, 2012; Zoghbi and Altenberg, 2018). Donor Tb^3+^-chelate fluorescence decays were recorded at each probe position in donor-only labeled MR^NBD^ complexes. Each Tb^3+^-chelate fluorescence decay fit well to two exponentials, with the longer lifetime comprising >85% of the signal. This longer lifetime was used in the distance analysis. From the Tb^3+^-chelate, Bodipy FL, and Cy3 lifetimes, the distances between donor and acceptor molecules (R) were calculated with

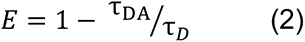

and

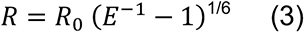

where E is the efficiency of energy transfer, τ_DA_ is the donor-sensitized lifetime of the acceptor (Bodipy FL or Cy3), τ_D_ is the lifetime of the donor (Tb^3+^), and R_0_ is the Förster distance between Tb^3+^ and Bodipy FL (44.9 Å) or Tb^3+^ and Cy3 (61.2 Å). Errors are the standard deviations from the mean of at least three measurements.

### HADDOCK

Molecular docking of the MR^NBD^ complex was done using the GURU interface on the HADDOCK 2.4 webserver (Dominguez et al., 2003; Van Zundert et al., 2016). For the three-body docking protocol, the PDB inputs were 3DSC (Pf Mre11 dimer lacking the helix-loop-helix (HLH) motifs and C-termini, DNAs deleted) (Williams et al., 2008) and two monomers of 3QKU (AMPPNP-bound Pf Rad50^NBD^ in complex with the Mre11 HLH motif) (Williams et al., 2011). Except for increasing the number of structures for rigid body docking (it0) to 3000, all settings used were default. The three linkers attaching the Pf Mre11 capping domain to the nuclease domain were allowed to be fully flexible (Y222-V236, Y249-G254, and V266-F273). To allow HADDOCK to move the two Rad50 monomers relative to each other, we input Rad50 as two identical monomers. C2 symmetry was enforced between the two Rad50^NBD^ monomers and between the two monomers of the Mre11 dimer.

HADDOCK Mre11 to Rad50 active Ambiguous Interaction Restraints (AIRs) were based on *M. jannaschii* (3AV0)(Lim et al., 2011) and *E. coli* (6S6V) (Käshammer et al., 2019) ATP-γ-S-bound MR structures. Passive AIRs (solvent accessible surface neighbors of active residues) were automatically defined by HADDOCK based on these active AIRs. Two extra passive AIRs were included on the “back side” of the Mre11 capping domain to allow for the extended structure seen in ATP-free *T. maritima* MR (PDB: 3QG5) (Lammens et al., 2011). Supplemental Fig. S5 shows the location of these AIRs on the structures of Mre11 and Rad50. The same set of active and passive AIRs were used in all of the HADDOCK runs with 50% random exclusion of AIRs in each structure calculation. The measured LRET distances were input as unambiguous restraints and defined as the Cβ-Cβ distance between the LRET-labeled residues, ±5 Å. For distance restraints greater than 75 Å, the bounds were increased to ±7 Å as the lifetime fits giving these distances were in a more error-prone region (i.e., the flat section) of the Tb^3+^-Cy3 LRET efficiency curve. The model with the lowest HADDOCK score in each run was considered as the best structure. To ensure that the unambiguous distance restraints obtained from one probe position were not dominating the structure calculations, HADDOCK runs were performed with systematic dropouts of all restraints calculated from a specific cysteine position (Supplemental Fig. S3). For the closed model all of the dropout structures maintained the proper D-loop/Walker A juxtaposition. Pymol version 2.4 was used to calculate RMSD values for all-atom alignments of HADDOCK models and to make figures of structures.

### SAXS and FoXS analysis

MR^NBD^ samples for SAXS were made by mixing Mre11 and Rad50^NBD^ in a 1:1.1 molar ratio. Samples were heated at 60 °C for 15 min to form complex and then cooled on the benchtop. The complex was then loaded onto a HiLoad 16/60 Superdex 200 column (GE Healthcare) equilibrated in 25 mM Hepes, 0.2 M NaCl, 0.1 mM EDTA, 1 mM TCEP, pH 7. The MR^NBD^ peak was collected and concentrated (Vivaspin, Sartorius) before dialysis (Slide-A-Lyzer MINI, Thermo Scientific) into LRET Buffer (50 mM Hepes, 100 mM NaCl, 5 mM MgCl_2_, 0.1 mM EDTA, 1% glycerol, 1 mM TCEP, pH 7) ± 2 mM ATP overnight at room temperature. Samples were diluted to 4 mg/mL (∼22 μM MR^NBD^ complex) with the equilibrated dialysis buffer and sent to the SIBYLS beamline 12.3.1 at the Advanced Light Source in Berkeley for high-throughput SAXS analysis (Classen et al., 2013; Dyer et al., 2014; Hura et al., 2009). For each sample, a total of 33 frames were collected at 0.3 sec intervals at 10 °C with a cell thickness of 1.5 mm. The sample to detector distance was 2 meters and the beam wavelength was 11 keV/1.27 Å. Buffer profiles were subtracted from sample profiles and the buffer subtracted frames for each sample were averaged using SAXS FrameSlice (https://sibyls.als.lbl.gov/ran).

Compared to the 3QKU PDB used as the input for HADDOCK runs, the Rad50^NBD^ protein construct used in the LRET and SAXS samples has an additional 7 amino acids on one side of the truncated coiled-coils and 8 amino acids on the other and the linker (GGAGGAGG) connecting them is slightly longer. To ensure that the structural models were as close to the protein construct used for the SAXS experiments as possible, the Rad50 coiled-coil and linker along with the 14 amino acid loop connecting the Mre11 helix-loop-helix and nuclease domain were modeled with loop modeling via Rosetta (version 3.11)(Mandell et al., 2009; Wang et al., 2007). This resulted in models only missing the 47 C-terminal amino acids of Mre11 (Supplemental Fig. S6). The FoXS server was used to back-calculate SAXS profiles of these modified closed, partially open, and open HADDOCK-Rosetta models and to fit these to the experimental MR^NBD^ scattering curves. The MultiFoXS server was used to calculate population-weighted ensemble fits(Schneidman-Duhovny et al., 2016, 2013).

## Acknowledgments

We thank Dr. Mariana Fiori and Prof. Guillermo Altenberg (TTUHSC, Lubbock, TX) for use of the fluorimeter for LRET data collection, technical help, and suggestions. We also thank Dr. Greg Hura (SIBYLS) for helpful suggestions. SAXS data was collected at SIBYLS which is supported by the DOE-BER IDAT DE-AC02-05CH11231 and NIGMS ALS-ENABLE (P30 GM124169 and S10OD018483). We also thank the National Institutes of Health (grant number: 1R35GM128906), CPRIT (grant number: RP180553), and The Welch Foundation (grant D-1876) for financial support to M.P.L.

## Author contributions

M.D.C. and M.P.L. conceived the project; M.D.C. performed the experiments and data analysis; M.D.C. and M.P.L. generated the visualizations; M.P.L. supervised the project; M.D.C. and M.P.L. wrote and edited the manuscript.

## Supplementary Materials for

**Fig. S1.**
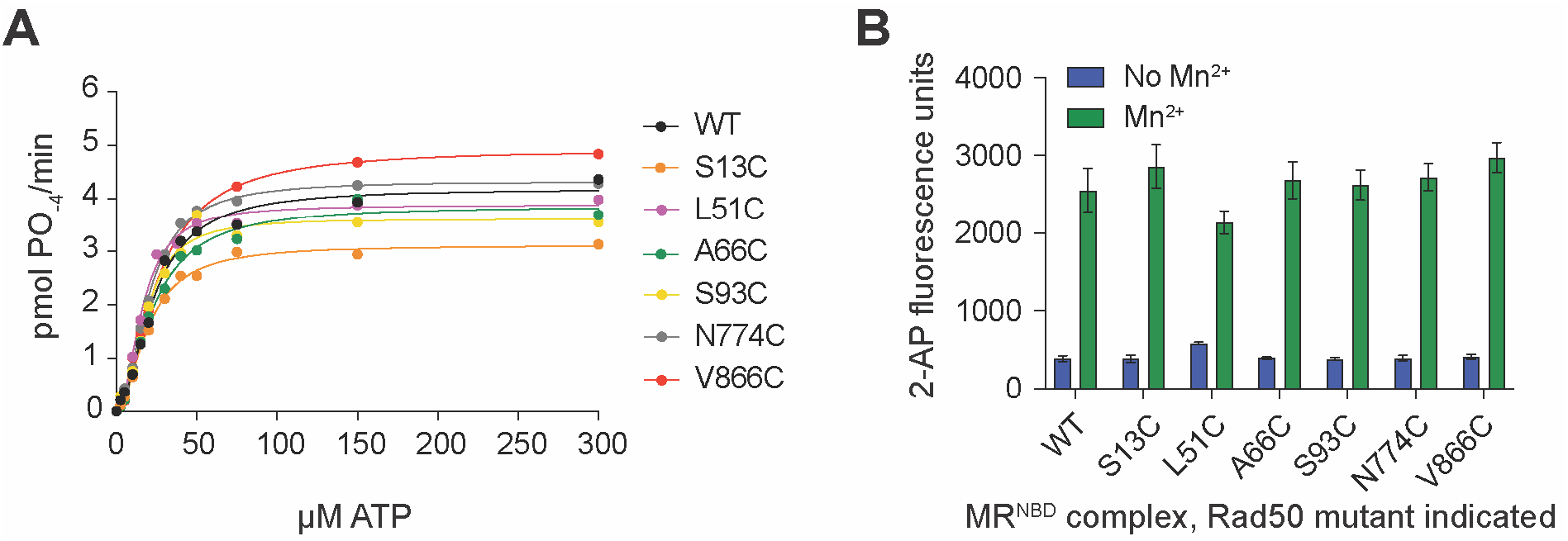
MR^NBD^ complexes made with cysteine mutants of Rad50 are active. (A) Steady-state Rad50 ATP hydrolysis kinetics for single cysteine mutants of MR^NBD^. Lines are the best fit to the Michaelis-Menten Hill equation. (B) Mre11 Mn^2+^-dependent exonuclease activity of single cysteine mutants of MR^NBD^ as determined by the Exo2 substrate in the absence (blue) and presence (green) of 1 mM MnCl_2_. Bars and errors represent the mean and standard deviation of at least three replicates.

**Fig. S2.**
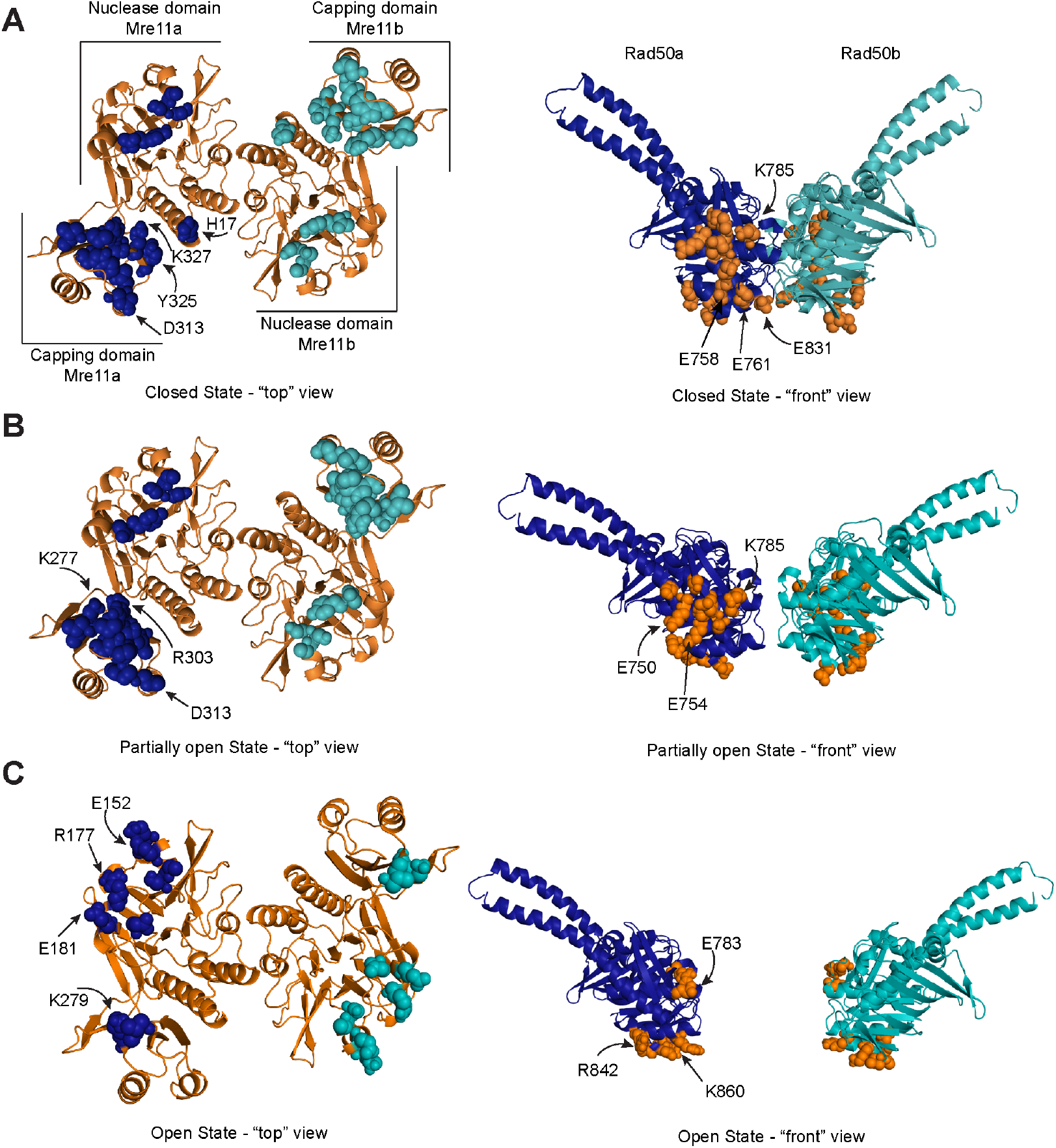
Mre11 and Rad50 make different interactions in the three conformations of the ATP-bound MR^NBD^ complex. (A) Mre11, left, and Rad50, right, residues within 3 Å of the other protein are highlighted as spheres on the closed HADDOCK model. Dark blue and teal spheres are Mre11 residues close to Rad50 and orange spheres are Rad50 residues close to Mre11. A number of residues mentioned in the text are indicated with arrows. (B) Mre11-Rad50 interactions in the partially open HADDOCK model. (C) Mre11-Rad50 interactions in the open HADDOCK model.

**Fig. S3.**
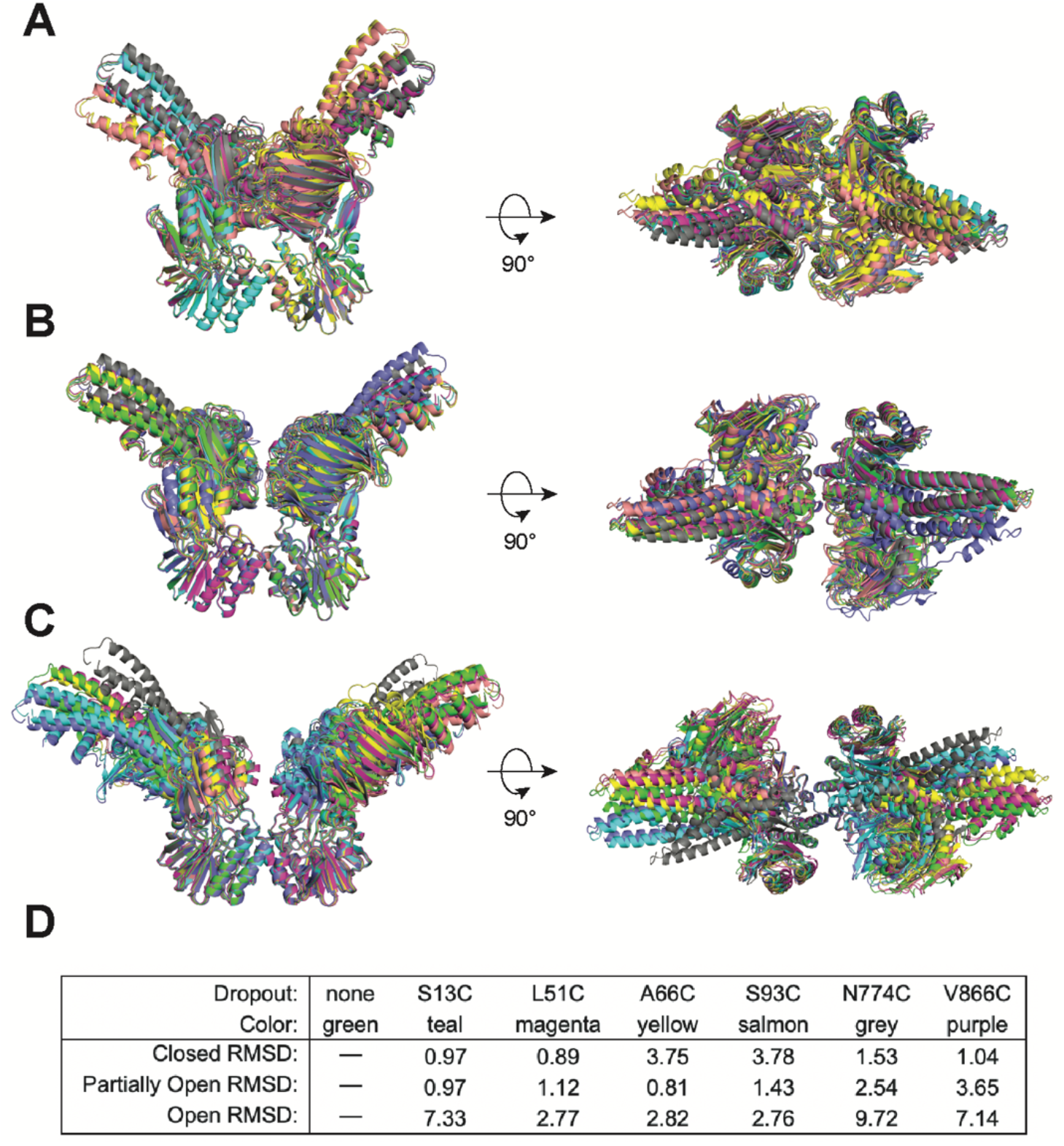
Data from one single LRET probe position do not dominate the HADDOCK structure calculations. HADDOCK models of (**A**) closed, (**B**) partially open, and (**C**) open conformations overlaid and aligned using Mre11. Each color is the model resulting from dropping all of the LRET unambiguous distance restraints associated with that specific probe position. (**D**) Table indicating the color for each dropout model in the overlays in A-C and the all-atom RMSD values (Å) calculated by Pymol for each when compared to the model using all of the LRET data for that conformation (i.e., no dropout).

**Fig. S4.**
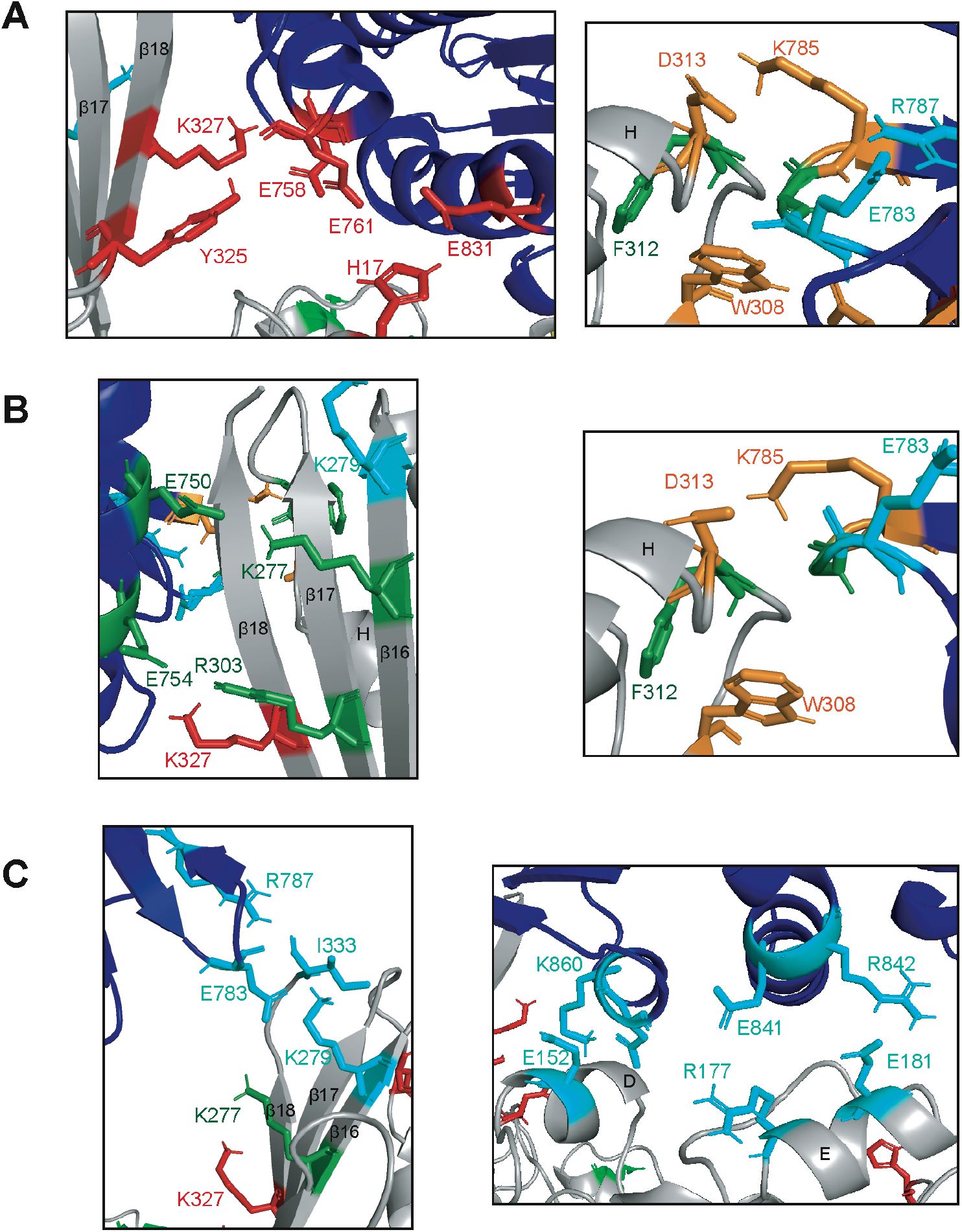
Unique interactions are made between Mre11 and Rad50 in the three conformations of MR^NBD^. (**A**) Specific interactions between residues in Rad50 (dark blue) and the Mre11 capping domain (grey) in the closed conformation. (**B**) Specific interactions between residues in Rad50 (dark blue) and the Mre11 capping domain (grey) in the partially open conformation. (C) Specific interactions between residues in Rad50 (dark blue) and the Mre11 capping domain (grey, left) or Mre11 nuclease domain (grey, right) in the open conformation. (**A-C**) Red residues interact only in closed. Orange residues interact in both closed and partially open. Green residues interact only in partially open. Blue residues interact only in open. A number of Mre11 β-sheets and α-helices are labeled.

**Fig. S5.**
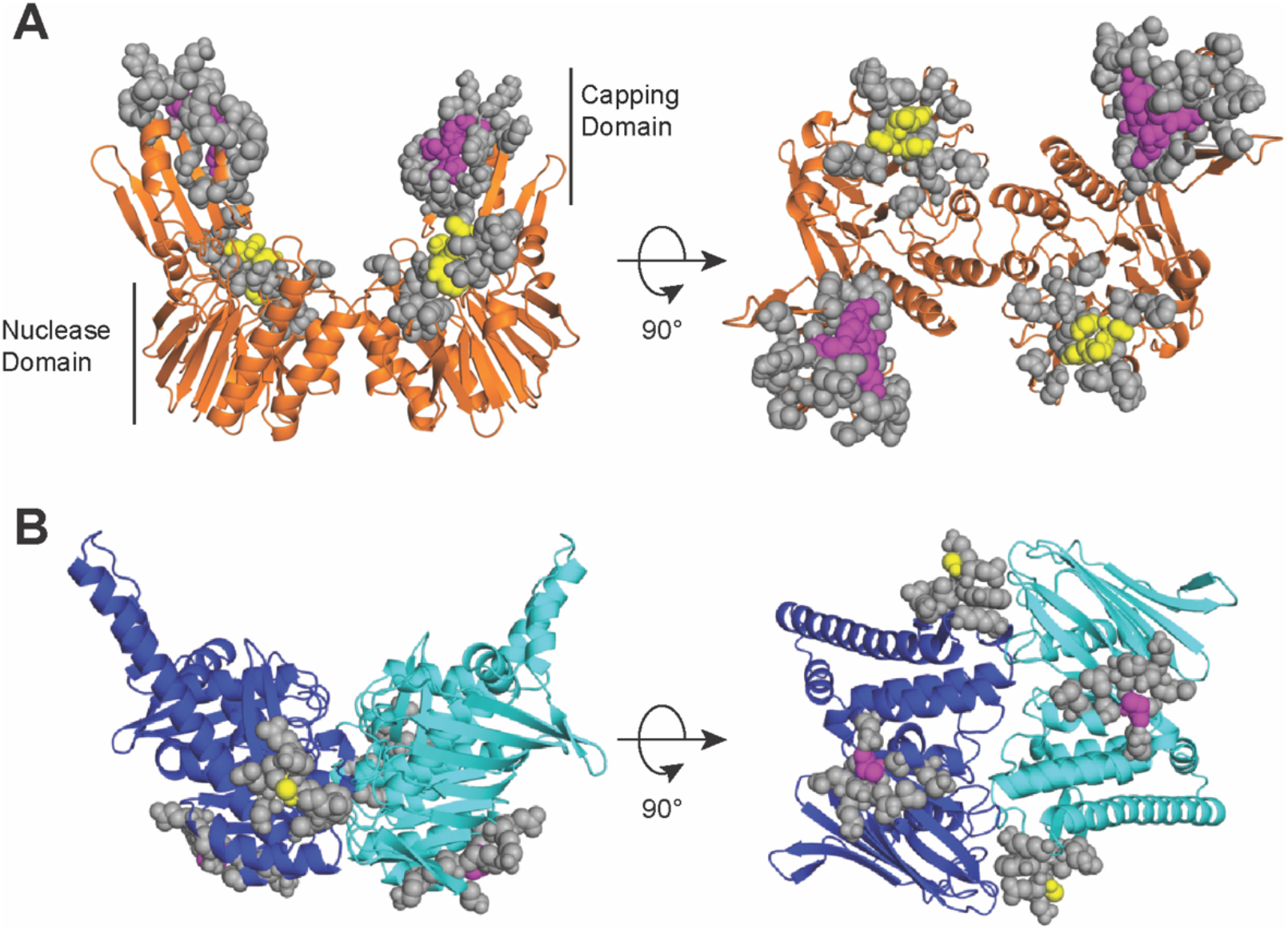
HADDOCK active and passive Mre11 to Rad50 Ambiguous Interaction Restraints (AIRs) are shown on the closed HADDOCK model. (**A**) Mre11 residues at the interface with Rad50 are indicated as spheres. Purple spheres in the capping domain (aa 308, 314, 328-330) and yellow spheres in nuclease domain (aa 147-150) were defined as active, while grey spheres were defined as passive. (**B**) Rad50 residues at the interface with Mre11 are indicated as spheres. Purple spheres (aa 864) and yellow spheres (aa 784) were defined as active, while grey spheres were defined as passive.

**Fig. S6.**
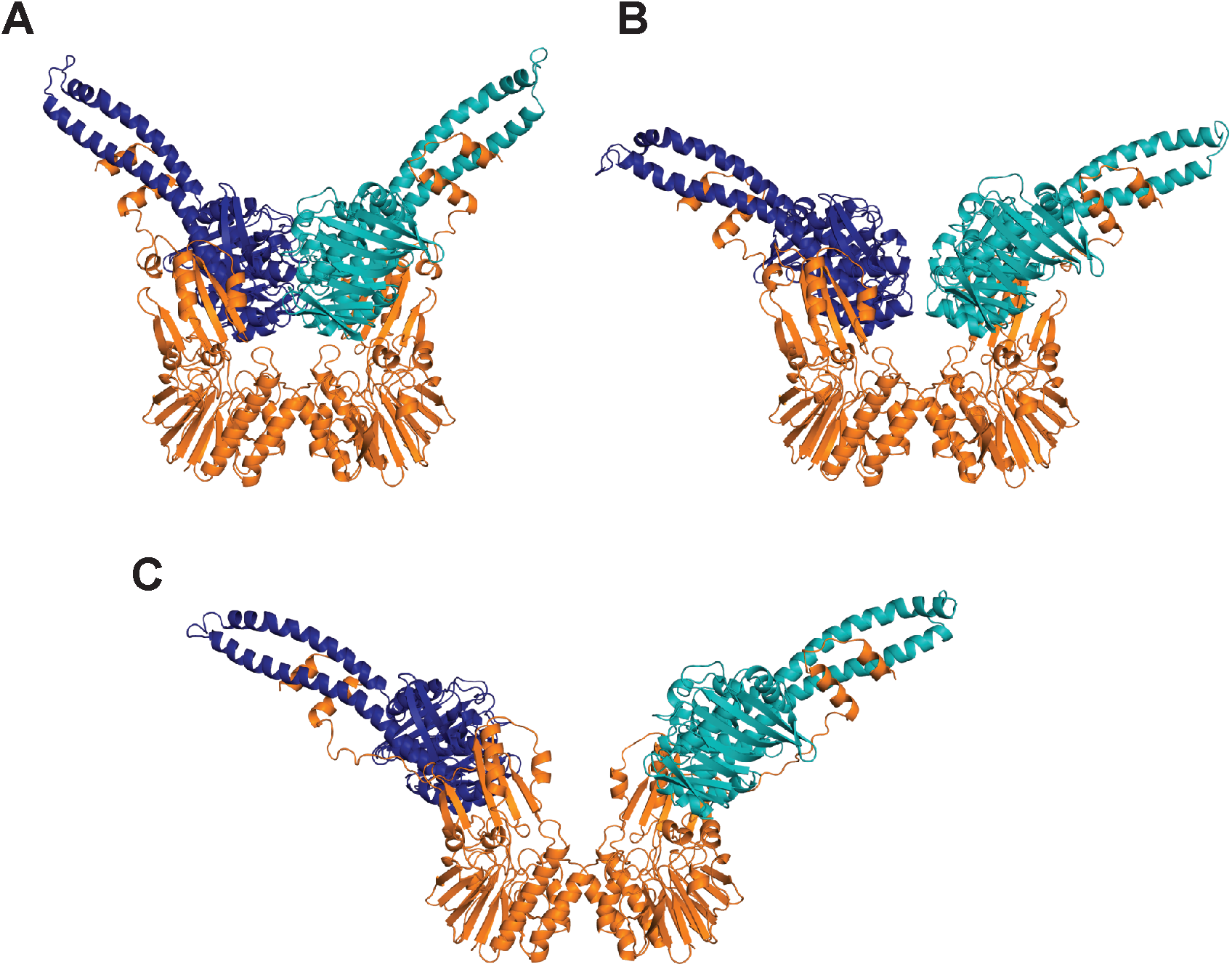
Rosetta-refined models of the MR^NBD^ conformations. Structures of (**A**) closed, (**B**) partially open, and (**C**) open conformations of MR^NBD^ where Rosetta was used to model in Mre11 residues 334-347 (the linker from the capping domain to the HLH motif) and to extend the Rad50 coiled-coils by 7 residues on one coil and 8 residues on the other, linking them with GGAGGAGG sequence, on the LRET-HADDOCK models. Mre11 is orange and the two Rad50 protomers are dark blue and teal. These structures were used in the FoXS and MultiFoXS analysis.

**Table S1.** MR^NBD^ LRET probe pair distances. Each row represents data from a unique LRET pair. Distances (in Å) were calculated from the decay of donor-sensitized Bodipy or Cy3 fluorescence emission as described in Methods. Errors are the standard deviation of n ≥ 3 LRET measurements.

| Bodipy position | Cy3 position | Tb<sup>3+</sup> position | 3QKU Cβ-Cβ distance | LRET Closed distance | LRET Partially open distance | LRET Open distance |
| --- | --- | --- | --- | --- | --- | --- |
| | S13C | S13C | 42.2 | | 50.9 ± 1.1 | 77.8 ± 1.7 |
| S13C | | L51C | 41.8 | 36.7 ± 1.1 | 51.4 ± 0.9 | |
| L51C | | S13C | 41.8 | 34.4 ± 1.1 | 51.2 ± 0.5 | |
| | S13C | L51C | 41.8 | | 51.6 ± 0.3 | 79.7 ± 3.0 |
| | L51C | S13C | 41.8 | | 51.3 ± 0.3 | 78.9 ± 1.9 |
| | A66C | S13C | 52.2 | 52.7 ± 1.2 | 75.4 ± 1.0 | |
| | S13C | S93C | 54.6 | 50.3 ± 0.7 | 76.8 ± 0.5 | |
| | S93C | S13C | 54.6 | 49.7 ± 1.7 | 79.5 ± 1.5 | |
| S13C | | N774C | 33.9 | 37.0 ± 0.9 | 48.7 ± 0.5 | |
| | S13C | N774C | 33.9 | | 44.9 ± 1.4 | 81.0 ± 0.5 |
| V866C | | S13C | 38.5 | 37.0 ± 0.1 | 49.2 ± 0.3 | |
| S13C | | V866C | 38.5 | 34.0 ± 0.7 | 50.2 ± 0.5 | |
| | S13C | V866C | 38.5 | | 48.7 ± 1.8 | 80.4 ± 3.3 |
| L51C | | L51C | 37.6 | 37.7 ± 0.6 | 45.4 ± 1.1 | |
| | L51C | L51C | 37.6 | | 47.0 ± 1.0 | 78.0 ± 2.0 |
| A66C | | L51C | 48.6 | 49.3 ± 1.3 | | |
| L51C | | A66C | 48.6 | 52.9 ± 1.7 | | |
| | A66C | L51C | 48.6 | | 56.3 ± 2.0 | 83.9 ± 4.1 |
| | L51C | A66C | 48.6 | | 55.9 ± 1.2 | 80.6 ± 3.9 |
| S93C | | L51C | 47.5 | 48.1 ± 0.8 | | |
| L51C | | S93C | 47.5 | 51.6 ± 0.4 | | |
| | S93C | L51C | 47.5 | | 55.1 ± 2.1 | 80.4 ± 3.4 |
| | L51C | S93C | 47.5 | | 53.1 ± 2.0 | 83.2 ± 1.1 |
| N774C | | L51C | 29.8 | 35.2 ± 0.6 | 50.3 ± 1.3 | |
| L51C | | N774C | 29.8 | 32.2 ± 1.0 | 50.6 ± 1.6 | |
| | L51C | N774C | 29.8 | | 47.1 ± 1.4 | 83.0 ± 0.6 |
| V866C | | L51C | 46.8 | 47.6 ± 0.7 | | |
| L51C | | V866C | 46.8 | 49.0 ± 0.3 | | |
| | V866C | L51C | 46.8 | | 52.4 ± 1.9 | 81.3 ± 2.5 |
| | L51C | V866C | 46.8 | | 52.2 ± 3.4 | 79.7 ± 1.7 |
| A66C | | A66C | 61.3 | 52.1 ± 1.5 | | |
| | A66C | A66C | 61.3 | 49.2 ± 1.3 | 81.5 ± 1.9 | |
| | A66C | S93C | 61.6 | 48.5 ± 2.1 | 78.7 ± 1.2 | |
| | S93C | A66C | 61.6 | 47.1 ± 1.8 | 82.1 ± 2.5 | |
| A66C | | V866C | 49.2 | 50.4 ± 1.3 | | |
| | V866C | A66C | 49.2 | | 55.0 ± 1.5 | 83.1 ± 2.0 |
| | A66C | V866C | 49.2 | | 53.7 ± 1.7 | 79.5 ± 2.4 |
| A66C | | N774C | 34.8 | 34.7 ± 0.6 | 49.3 ± 1.2 | |
| | A66C | N774C | 34.8 | | 47.2 ± 0.5 | 82.8 ± 1.3 |
| S93C | | S93C | 58.8 | 51.6 ± 1.2 | | |
| | S93C | S93C | 58.8 | $51.4 \pm 1.9$ | $79.9 \pm 0.6$ | |
| S93C | | N774C | 31.3 | $32.6 \pm 0.6$ | $49.9 \pm 1.0$ | |
| N774C | | S93C | 31.3 | $34.2 \pm 1.1$ | $50.6 \pm 0.6$ | |
| | N774C | S93C | 31.3 | | $45.6 \pm 1.0$ | $75.6 \pm 1.2$ |
| | V866C | S93C | 56.5 | $52.4 \pm 1.5$ | $80.1 \pm 1.0$ | |
| | S93C | V866C | 56.5 | $48.3 \pm 2.7$ | $78.8 \pm 2.7$ | |
| N774C | | N774C | 36.7 | $35.7 \pm 0.2$ | $51.5 \pm 2.3$ | |
| | N774C | N774C | 36.7 | | $45.8 \pm 1.8$ | $83.5 \pm 2.2$ |
| V866C | | V866C | 37.0 | $36.2 \pm 1.5$ | $53.5 \pm 0.8$ | |
| | V866C | V866C | 37.0 | | $50.6 \pm 1.8$ | $82.7 \pm 3.3$ |

**Table S2.** MR^NBD^ LRET experimental probe distances and HADDOCK model distances.

| Pair of Rad50 residues | 3QKU dimer Cβ-Cβ distance (Å) | LRET unambiguous restraint (Å) | | | HADDOCK model Cβ-Cβ distance (Å) | | |
| --- | --- | --- | --- | --- | --- | --- | --- |
|  |  | Closed (±5) | Partially open (±5) | Open (±7) | Closed | Partially open | Open |
| S13-S13 | 42.2 | 35.4 | 51.0 | 77.8 | 40.4 | 56.2 | 74.9 |
| L51-L51 | 37.6 | 37.7 | 46.2 | 78.0 | 36.1 | 51.6 | 82.0 |
| A66-A66 | 61.3 | 50.7 | 81.5 |  | 58.1 | 74.4 | 93.0 |
| S93-S93 | 58.8 | 51.5 | 79.9 |  | 56.2 | 73.4 | 97.0 |
| N774-N774 | 36.7 | 35.7 | 48.7 | 83.5 | 38.4 | 50.0 | 78.3 |
| V866-V866 | 37.0 | 36.2 | 52.1 | 82.7 | 36.1 | 46.9 | 73.4 |
| S13-L51 | 41.8 | 35.6 | 51.4 | 79.3 | 40.0 | 53.6 | 79.1 |
| S13-A66 | 52.2 | 52.7 | 75.4 |  | 49.5 | 65.7 | 84.0 |
| S13-S93 | 54.6 | 50.0 | 78.2 |  | 52.3 | 67.4 | 88.1 |
| S13-N774 | 33.9 | 37.0 | 46.8 | 81.0 | 32.9 | 41.8 | 73.2 |
| S13-V866 | 38.5 | 35.5 | 49.4 | 80.4 | 37.1 | 52.8 | 74.0 |
| L51-A66 | 48.6 | 51.1 | 56.1 | 82.3 | 46.4 | 61.2 | 86.2 |
| L51-S93 | 47.5 | 47.6 | 53.0 | 81.8 | 45.4 | 62.3 | 88.8 |
| L51-N774 | 29.8 | 33.7 | 49.3 | 83.0 | 29.2 | 44.4 | 77.1 |
| L51-V866 | 46.8 | 48.3 | 52.3 | 80.5 | 46.0 | 57.1 | 82.7 |
| A66-S93 | 61.6 | 47.8 | 80.4 |  | 58.8 | 74.7 | 95.8 |
| A66-N774 | 34.8 | 34.7 | 48.3 | 82.8 | 32.9 | 46.7 | 77.5 |
| A66-V866 | 49.2 | 50.4 | 54.3 | 81.3 | 47.3 | 63.4 | 83.1 |
| S93-N774 | 31.3 | 33.4 | 48.7 | 75.6 | 29.4 | 49.2 | 79.8 |
| S93-V866 | 56.5 | 50.4 | 79.5 |  | 54.9 | 69.7 | 90.0 |

**Movie S1.** The movie depicts the transitions between the closed, partially open, and open conformations first shown from the ‘side’ view and then from the ‘top’ view. Next, Mre11 is hidden to highlight the position of the Rad50 Walker A (N32, magenta), signature motif (S793, yellow), and D-loop (D829) in these three conformations. Mre11 is colored orange, whereas Rad50 is colored blue and teal. The morph between the conformations was generated in Chimera (version 1.15) and rendered in PyMOL (version 2.4).

